# Dentate gyrus morphogenesis is regulated by β-CATENIN function in hem-derived fimbrial glia

**DOI:** 10.1101/2022.09.15.508086

**Authors:** Arpan Parichha, Debarpita Datta, Varun Suresh, Mallika Chatterjee, Michael J. Holtzman, Shubha Tole

## Abstract

The dentate gyrus, a gateway for input to the hippocampal formation, arises from progenitors in the medial telencephalic neuroepithelium adjacent to the cortical hem. Dentate progenitors navigate a complex migratory path guided by two cell populations that arise from the hem, the fimbrial glia, and Cajal-Retzius (CR) cells. Since the hem expresses multiple Wnt genes, we examined whether β-CATENIN, which mediates canonical Wnt signaling and also participates in cell adhesion, is necessary for the development of hem-derived lineages. We report that the fimbrial glial scaffold is disorganized and CR cells are mispositioned upon hem-specific disruption of β-CATENIN. Consequently, the dentate migratory stream is severely affected, and the dentate gyrus fails to form. Using selective Cre drivers, we further determined that β-CATENIN function is required in the fimbrial glial scaffold, but not in the CR cells, for guiding the dentate migration. Our findings highlight a primary requirement for β-CATENIN for the organization of the fimbrial scaffold and a secondary role for this factor in dentate gyrus morphogenesis.

## Introduction

In the developing central nervous system, specialized cell types arising from discrete neuroepithelial domains perform important roles in guiding the assembly of particular brain structures. The cortical hem, a Wnt-rich structure in the embryonic medial telencephalon, produces several distinct lineages, including the telencephalic choroid plexus epithelium, Cajal-Retzius (CR) cells, and glia that comprise the fimbrial scaffold (Louvi et al., 2007, Gu et al., 2011). Each of these lineages contributes either secreted factors or cellular substrates that perform important functions in neocortical and hippocampal development (Caramello et al., 2021, Bagri et al., 2002, Lehtinen et al., 2011). The molecular processes that operate within hem progenitors to modulate the development of these lineages are not well understood. We focused on β-CATENIN because it performs two distinct functions, as a transcription factor in the canonical Wnt pathway, and also as part of the adherens complex at the cell membrane. Here, we examine the effect of loss of β-CATENIN on the development of the CR cells and the fimbrial glial scaffold.

Both, the CR cells and the fimbrial glial scaffold, are critical in the regulation of dentate gyrus morphogenesis, a process that has been well characterized in rodents. Dentate gyrus granule neurons are produced in the dentate neuroepithelium (DNE), a region of the ventricular zone adjacent to the hem (Altman et al., 1990a). Proliferating progenitors from the DNE as well as specified dentate granule cells migrate along a curved path called the dentate migratory stream (DMS; Altman et al., 1990b), and eventually form the characteristic “V” shaped gyrus at the hippocampal fissure. CR cells secrete chemokines that are critical for the guidance of the DMS (Nakajima et al., 1997, Bagri et al., 2002; Lu et al., 2002). The migration of the CR cells is itself dependent on the fimbrial glial scaffold (Gu et al., 2011). This scaffold is critical in guiding both the CR cells and the DMS, therefore, it is important to understand how its organization and orientation are regulated. Factors that regulate the timing of formation (Caramello et al., 2021) or the extent of this scaffold (Barry et al., 2008) have been reported, however, mechanisms that control its organization have not yet been identified.

Here, we report that when β-CATENIN is lost in embryonic mouse hem, the resulting fimbrial glial scaffold is specified, but is highly disorganized and protrudes ectopically into the ventricle. CR cells are specified, but mispositioned in the ectopic protrusion formed by the fimbrial glia. Dentate granule cells are also specified but their migration is profoundly disrupted resulting in the absence of a morphologically distinct dentate gyrus. These deficits do not appear if β-CATENIN is disrupted in the CR cells alone. Therefore, β-CATENIN function in the hem-derived fimbrial glial scaffold is necessary for its organization, and consequently for the morphogenesis of the dentate gyrus.

## Results and Discussion

### Lmx1aCre identifies hem-derived cell types

Lmx1a expression was seen in the E12.5 cortical hem and the choroid plexus (Fig. 1A). An Ai9 reporter (a line carrying a stop-floxed tdTomato cassette) driven by Lmx1aCre (Chizhikov et al., 2010) labeled the hem and its derivatives including BLBP+ fimbrial glia and REELIN+ CR cells at E12.5 and E18.5 (Fig. 1B-D). Dentate granule cells, identified by PROX1 labeling, do not display Ai9 fluorescence since they are not derived from the hem (Fig. 1C, D; Caramello et al., 2021).

**Figure 1:**
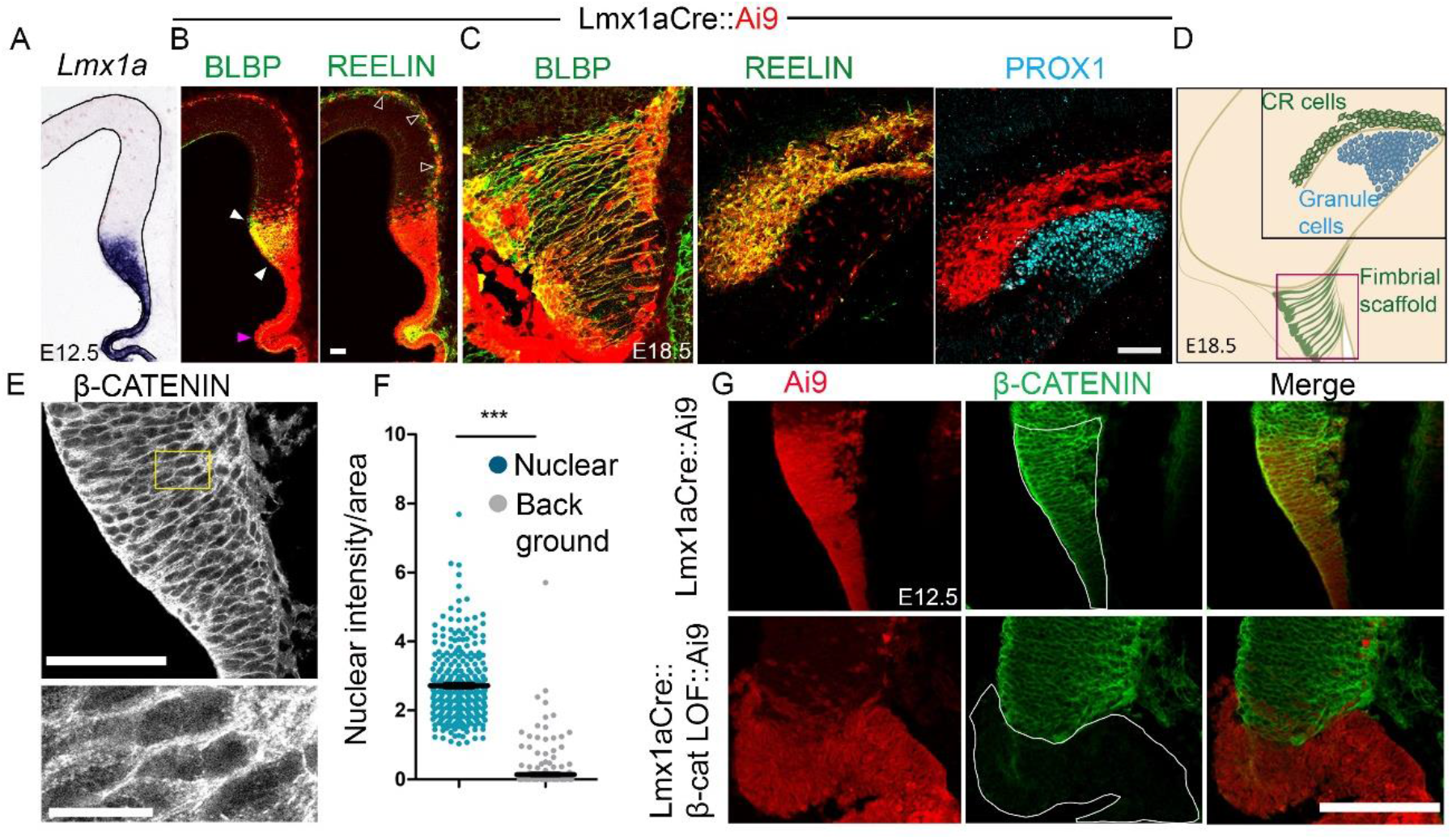
Lmx1aCre identifies hem-derived lineages. (A) At E12.5 Lmx1a expression is seen in the hem and the choroid plexus. (B) Lmx1aCre drives Ai9 expression in the BLBP+ fimbrial glia (white arrowheads), the REELIN+ CR cells (open arrowheads) and the choroid plexus epithelium (Magenta arrowhead). (C) At E18.5 Ai9 fluorescence co-localizes with Brain Lipid Binding Protein (BLBP) and REELIN but not with dentate granule cell marker PROX1. (D) Cartoon illustrating the hem-derived lineages and dentate granule cells at E18.5. The black box in D indicates the region displayed in the BLBP and PROX1 images in C and the magenta box represents the region displayed in the BLBP image. Representative images are shown of sections taken from N= 5 brains (biologically independent replicates) examined over 5 independent experiments. (E, F) Immunostaining and quantification of nuclear β-CATENIN (G) β-CATENIN is lost in Ai9 expressing cells in the Lmx1aCre::β-Catenin LOF brain (white outline). Scale bars: 10μm (E high mag), 50 μm (B, C, E and G).

### β-CATENIN loss of function (LOF) in the hem disrupts the fimbrial glial scaffold and perturbs CR cell migration

*β-Catenin (Ctnnb1)* is expressed throughout the telencephalic midline neuroepithelium from E11.5 (Parichha et al., 2022, Kadowaki et al., 2007). A scRNA-seq study of E14.5 mouse cortex also identified *β-Catenin* to be expressed in hem progenitors, CR cells, and choroid plexus epithelium (Loo et al., 2019, <u>GSE123335</u>). We disrupted *β-Catenin* in the developing hem using the Lmx1aCre driver together with a well-described mouse line in which exons 2-6 of the *β-Catenin (Ctnnb1)* gene are flanked by loxP sites and Cre mediated recombination resulted in a non-functional, transcriptionally inactive protein (Brault et al., 2001). In E12.5 Lmx1aCre::β-Catenin LOF brains, the BLBP-positive fimbrial glial scaffold was no longer confined to the region of the hem as it was in the controls (Fig. 2B). Instead, BLBP+ cells formed an ectopic protrusion into the ventricle just above the choroid plexus (Fig. 2B-C, E). We examined whether this protrusion arose as a result of excessive proliferation of the β-Catenin LOF hem progenitors. However, both PHH3 and Ki67 immunostaining revealed no apparent change in proliferation (Fig. S1F-I)

**Figure 2:**
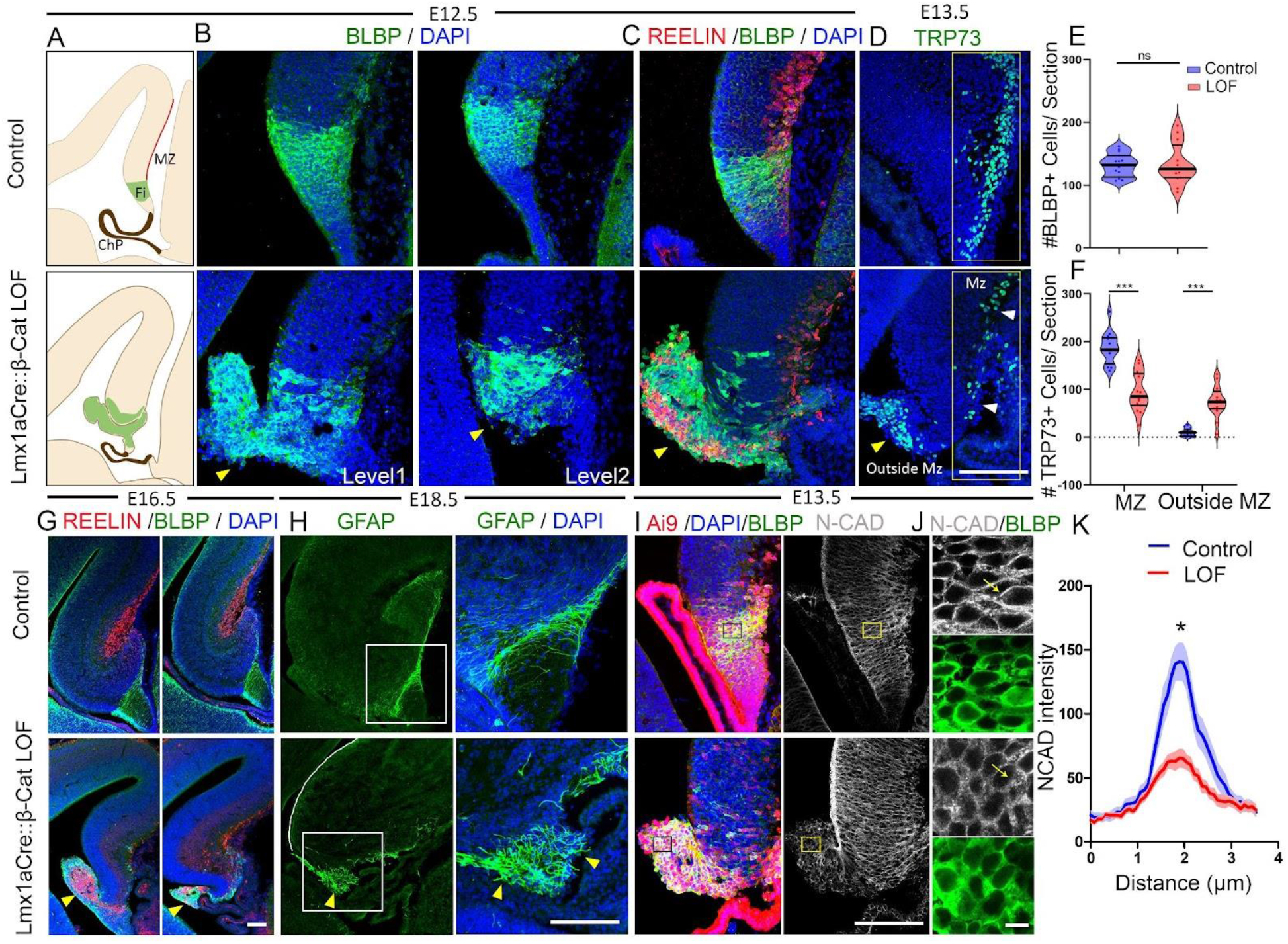
Loss of β-CATENIN in the cortical hem causes disrupted assembly of the fimbrial scaffold and mispositioning of the CR cells. (A) Cartoons of E12.5 sections indicating the marginal zone (red line, MZ) and the fimbria (green, Fi) (B) BLBP immunostaining at two levels of sectioning reveals the disorganization of fimbrial scaffold and an ectopic glial protrusion (yellow arrowhead) in Lmx1aCre::β-Catenin LOF brains in comparison with controls. Representative images are shown of sections taken from N=5 brains (biologically independent replicates) examined over 5 independent experiments. (C) Co-immunolabeling for REELIN and BLBP reveals the mispositioning of CR cells in the ectopic glial protrusion (yellow arrowhead); representative images are shown of sections taken from N=3 brains (biologically independent replicates) examined over 2 independent experiments. (D) At E13.5, hem-derived TRP73+ CR cells are normally localized to the marginal zone, (white arrowheads) but this population is significantly decreased in Lmx1aCre::β-Catenin LOF brains, and instead, the cells are mislocalized in the ectopic glial protrusion (yellow arrowhead). (E) Quantification of BLBP+ cells shown in (B) reveals no significant difference in total BLBP+ cell number. (F) Quantification of TRP73+ cells shown in (D) indicates that 95% of the CR cells are localized in the MZ region in control brains, whereas only 56% are localized in the MZ in Lmx1aCre::β-Catenin LOF brains and 44% are found outside the MZ region. N=5 brains (biologically independent replicates) examined over 4 independent experiments. (G) Co-immunolabeling with BLBP and REELIN reveals the defective fimbrial scaffold in Lmx1aCre::β-Catenin LOF brains at E16.5. (H) GFAP labeling at E18.5 reveals a well-ordered fimbrial scaffold in controls, but a disorganized scaffold in the Lmx1aCre::β-Catenin LOF brains. (I, J) Fimbrial scaffold cells identified by BLBP immunolabeling also display N-CADHERIN. (J) Images of the boxed region in (I) and quantification (K) of N-CADHERIN across cell junctions along the yellow arrow (J), n=30 cells from N=3 brains (biologically independent replicates) examined over 3 independent experiments. Statistical tests (E, F, & K): Two-tailed unpaired Mann Whitney U test, * p < 0.05, ** p < 0.01, *** p < 0.001, ns if p-value> 0.05; P=0.9268 (E), P=0.007937 (F), P=0.0150 (K). Scale bars: 10 μm (I high mag); 100 μm (all panels in B-I).

One established function of the glial scaffold is to provide critical cellular guidance to hem-derived CR cells in their migration to the marginal zone (MZ) of the hippocampus (Gu et al., 2011). REELIN immunostaining revealed many of these cells to be accumulated in the ectopically protruding fimbrial mass in Lmx1aCre::β-Catenin LOF brains (Fig. 2C). We used a specific hem-derived CR cell marker, TRP73 to quantify the mislocalization of the CR cells (Fig. 2D). In E13.5 control brains, 95% of the CR cells occupied the MZ region and 5% were found outside this region, presumably en route to their final destination. Upon loss of β-CATENIN however, 56% were found in the MZ region and 43% were found outside the MZ (Fig. 2D, 2F).

The fimbrial glial scaffold dysmorphia seen in E12.5 β-Catenin LOF brains did not improve with time but persisted at later stages in development (Fig. S2A). A comprehensive time-course panel from E11.5 to postnatal day (P)2 reveals misoriented fibers revealed by Glial Fibrillary Acidic Protein (GFAP) or BLBP labeling, in contrast to the well-organized alignment of these fibers in control brains (Fig. 2H; Fig. S2A). REELIN-positive cells were mislocalized within the protruded glial mass at all stages examined (Fig. 2G; Fig S2B).

In summary, loss of β-CATENIN in the hem resulted in striking defects in the organization of the fimbrial glial scaffold and in the migration of the CR cells, although both cell types appeared to be specified and displayed appropriate markers.

### Transcriptional and Adhesion-related roles of β-CATENIN in the hem

It is well established that β-CATENIN has two roles: it mediates transcriptional regulation in the canonical Wnt signaling pathway (Valenta et al., 2012) and also participates in adherens junctions where it bridges the CADHERINS with the ACTIN network (Gumbiner, 2005). Deletion of exon 2-6 of the *β-Catenin* gene disrupts both functions (Valenta et al., 2011). No detectable β-CATENIN is seen in the Ai9-expressing cells in Lmx1aCre::β-Catenin LOF brains (Fig.1G). We performed RNASeq analysis of the E12.5 control and β-Catenin LOF hem. Gene ontology analysis shows GO:BP terms that include both “Canonical Wnt signaling pathway” and “Positive regulation of cell junction assembly” to be significantly downregulated upon loss of β-CATENIN, indicating that both functions of β-CATENIN may be relevant for fimbrial scaffold morphogenesis (Fig. S3C).

Several members of the canonical Wnt signaling pathway including *Fzd1, Tcf7l1, Tcf7l2, Axin2*, and nuclear LEF1 were present in the E12.5 hem (Fig. S3E-G). Canonical Wnt targets *Pcsk6, Fzd10, Runx2, Sost, Axin2, Wnt3* are significantly downregulated in the transcriptome of the β-Catenin LOF hem (Fig. S3H, Supplementary table 1). We further validated that canonical Wnt targets *Axin2* and LEF1 are down regulated in β-Catenin LOF hem by using in situ hybridization and immunohistochemistry respectively (Fig. S3I-J). Finally, canonical Wnt signaling was implicated in the development of a different glial scaffold located at the dentate gyrus, which displayed defects upon loss of Wnt co-receptor LRP6 and transcriptional mediator LEF1 (Zhou et al., 2004). Together, these findings suggested that the fimbrial glial scaffold may require canonical Wnt signaling for normal development.

In contrast to the dysregulation of canonical Wnt targets, the mRNA and protein levels for fimbrial markers like GFAP, FABP7, ALDH1L1 and AQP4 did not differ significantly (Fig. S4A and B). Transcription factors SOX9, NFIA, and NFIB required for normal development of the fimbrial glial scaffold (Caramello et al., 2021; Bunt et al., 2017) are apparently unaltered in expression levels in upon loss of β-CATENIN (Fig. S4C and D).

To investigate the role of β-CATENIN in adhesion, we examined the distribution of N-CADHERIN, a key member of cell-cell adherens junctions in neuroepithelium (Hirano and Takeichi, 2012), and enriched at the embryonic telencephalic midline (Kadowaki et al., 2007). We focused on the cell boundaries to assess the junctional integrity of the control and mutant hem (E13.5, Fig. 2 I-K) and fimbrial glial cells (E16.5, Fig. S5). Hem and fimbrial glial cells were identified by co-immunolabeling for BLBP. In the control, N-CADHERIN distribution measured along a linear path traversing two neighboring cells reveals a sharp peak in intensity at the boundary between the cells. In β-Catenin LOF brains, there was a marked flattening of the peak intensity at cell boundaries, indicating a diffused distribution of N-CADHERIN, and suggesting altered cell-cell adhesion, although the mRNA levels of *Cdh2*, which encodes N-CADHERIN, appear unaltered (Fig. 2K; Fig. S5F and G).

These results suggested that the transcriptional dysregulation of factors that control fimbrial scaffold morphogenesis and/or perturbed cell adhesion as a result of altered N-CADHERIN distribution may underlie the disruption of the fimbrial glial scaffold resulting in the ectopic protrusion seen in β-Catenin LOF brains. A comparison with β-Catenin LOF selective to its transcriptional role, leaving its cell adhesion intact (Valenta et al., 2011), would be a useful additional study in this context and may help to identify novel β-CATENIN targets responsible for proper orientation and positioning of the fimbrial scaffold.

### The dentate gyrus migratory stream is misdirected upon loss of β-CATENIN in the cortical hem which leads to malformation of the dentate gyrus

That the fimbrial glial scaffold is specified but disorganized in Lmx1aCre::β-Catenin LOF brains offered an opportunity to examine the effects of this perturbation on dentate morphogenesis. Dentate gyrus granule neurons are produced in the DNE and express PROX1, a factor necessary for their maturation (Lavado et al., 2010). Proliferating progenitors (SOX9+, PAX6+) from the DNE are guided by and migrate together with TBR2+ intermediate progenitors along with the DMS, to the final position of the dentate gyrus, where proliferation continues (Fig. 3A; Nelson et al., 2020). The DNE, the migration path, and the final destination are termed the primary, secondary, and tertiary matrices respectively (Altman et al., 1990b, Sugiyama et al., 2013).

**Figure 3:**
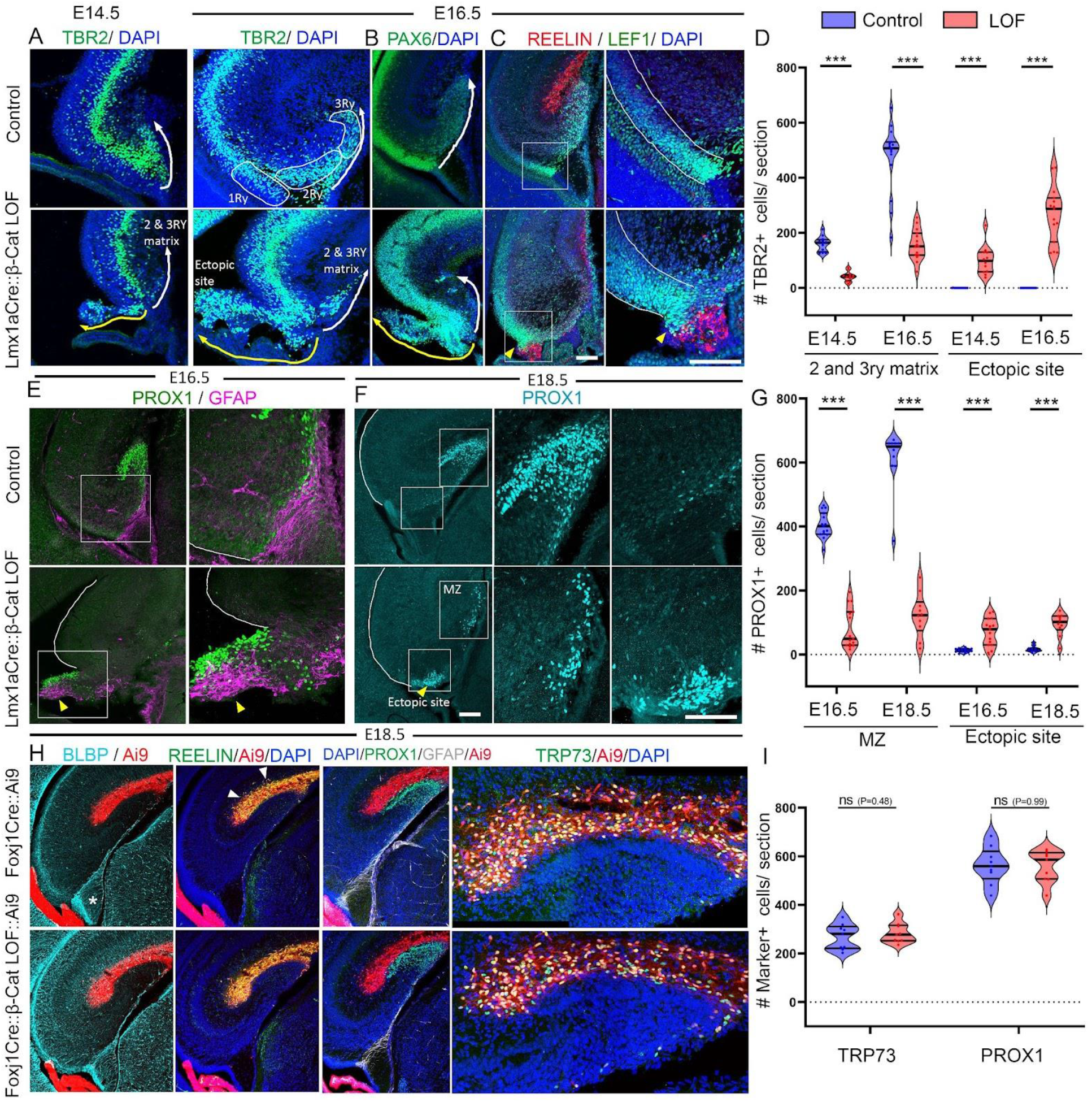
Loss of β-CATENIN in cortical hem leads to mis-migration of the dentate migratory stream resulting in defective DG morphogenesis. (A, B) TBR2+ and PAX6+ progenitors are seen in the 2 and 3ry matrix of the DMS at E14.5 and E16.5, in control brains (DMS, white arrow). In Lmx1aCre::β-Catenin LOF brains, these cells are diverted to the ectopic glial protrusion (yellow arrow); representative images are shown of sections taken from N=4 (E14.5), N=5 (E16.5) brains (biologically independent replicates) examined over 3 independent experiments. (C) In control brains, REELIN+ CR cells reside in the hippocampal fissure above LEF1+ DMS cells, whereas in β-Catenin LOF brains both these cell types are co-mingled in the ectopic protrusion (yellow arrowhead). (D) Quantification of the images in (A). (E-G) PROX1+ dentate granule cells localize to the dentate gyrus in controls, but are diverted to the ectopic protrusion in β-Catenin LOF brains (yellow arrowhead). The boxed region in C, E and F are shown in high magnification in the adjacent images. (H) FoxJ1Cre does not drive Ai9 reporter expression in the BLBP+ fimbrial scaffold (asterisk) but efficiently labels the REELIN+ CR cells (white arrowheads). At E18.5, PROX1+ dentate granule cells and TRP73+ CR cells appear normally positioned in both control and Foxj1Cre::β-Catenin LOF brains (I) Quantifications of the images in (H). D: N= 4 (E14.5); N=5 (E16.5); G: N=5 (E16.5 and E18.5); I: N=3 (E18.5) biologically independent replicates examined over 3 independent experiments. In Fig. 3A and H, image stitching was performed. Statistical tests (D, G & I): Two-tailed unpaired Mann Whitney U test, *p < 0.05, **p < 0.01, ***p < 0.001, ns if p-value> 0.05; P=0.9268 (E), P=0.000011 (G, E16.5 and E18.5 MZ), P=0.000084 (G, E16.5 ectopic site), P=0.000076 (G, E18.5 ectopic site), P=0.489428 (I, TRP73), and P>0.999999 (I, PROX1). All scale bars: 100 μm.

Upon loss of β-CATENIN in the hem, both TBR2+ and PAX6+ progenitors were found to accumulate in the ectopic protrusion that contained the fimbrial glia and Cajal-Reizius cells, instead of migrating along with the DMS (Fig. 3A, B). LEF1, a factor necessary for dentate granule fate (Galceran et al., 2000), marks a population in the DNE which migrates to the tertiary matrix in controls. In Lmx1aCre::β-Catenin LOF brains, LEF1+ cells accumulated close to the ectopic CR cell cluster, and very few migrated to their final destination at the hippocampal fissure (Fig. 3C). Finally, PROX1+ dentate granule cells failed to form the characteristic “V” shape of the dentate gyrus in β-Catenin LOF brains, and appear arrested and juxtaposed to the ectopic and disorganized fimbrial glial protrusion (Fig. 3E, F). We quantified the percentage of TBR2+ and PROX1+ cells that reached the tertiary matrix/ dentate gyrus or were mislocalized in the region of the hem/ ventricular zone, and found that whereas the control brains had a negligible number of mislocalized cells, β-Catenin LOF brains contained a significantly larger number in the ectopic site, and a significantly smaller number in the location of the future dentate gyrus (Fig. 3D, G, Fig. S6 A-F). The TBR2+ intermediate progenitor population in the 2ry and 3ry matrix of β-Catenin LOF brains was similar to that in controls (Figure S6D. However, the total number of PROX1 cells in the β-Catenin LOF brain was significantly less than that in controls (Figure 3G; Supplementary Figure S6H), suggesting that reduced numbers of DG progenitors were specified, or their proliferation/ survival was affected. DG specification requires Wnt signaling from the hem (Lee et al., 2000). While it appears that DG cell fate was specified at least in terms of PROX1, the underlying basis of the reduction in PROX1+ cells is an important angle for further studies focused on the mechanism of specification of DG cells.

Both CR cells and the fimbrial scaffold glia are important in guiding the DMS (Stanfield, B.B. and Cowan, W.M. 1979, Caramello et al., 2021). When chemokine signaling from CR cells is disrupted in either CXCR4 or CXCL12/SDF-1 mutants (Bagri et al., 2002; Lu et al., 2002), the DMS is perturbed. The CR cells are themselves guided by the fimbrial glial scaffold. When this scaffold ablated from E13.5 using DTA driven by hem-specific Fzd10Cre, CR cell migration is aberrant, and these cells are scattered in ectopic locations (Gu et al., 2010). To test the requirement for β-CATENIN specifically in the CR cells, we used Foxj1Cre which is selective for these cells and does not drive recombination in the fimbrial scaffold (Fig. 3H). Foxj1Cre::β-Catenin LOF brains showed no defect in CR cell localization in the hippocampal fissure and there is no apparent defect in the formation of the dentate gyrus (Fig. 3H, I).

In summary, we have identified that β-CATENIN function is critically required in the hem-derived fimbrial glial scaffold (but not in hem-derived CR cells), for normal morphogenesis of the dentate gyrus. In the absence of β-CATENIN, the fimbrial glial scaffold is disorganized and forms an ectopic ventricular protrusion as early as E12.5, prior to the migration of Prox1+ dentate granule cells which begins at E14.5. Both, dentate granule cells and CR cells depend on the glial scaffold for their proper migration and both these cell types migrate into the protrusion instead (Fig. 4, Fig. S7, Supplementary movie 1). This disrupted migration does not improve with time, and the dentate gyrus fails to form. Therefore, the protrusion which is composed of disorganized glial scaffold cells in Lmx1aCre::β-Catenin LOF brains is the major and proximate cause of the profoundly defective morphogenesis of the dentate gyrus. While deficits in the organization of glia in the dentate region have been reported in some canonical Wnt pathway mutants such Lrp6 and Lef1 (Zhou et al., 2004), neither of these, nor any known mutant that lacks a component of adherens junctions (Zhang et al., 2013), displays such extreme disorganization as that seen in Lmx1aCre::β-Catenin LOF brains. These results highlight the function of a single factor, β-CATENIN, as a key regulator of fimbrial scaffold organization, which ultimately controls the morphogenesis of a key hippocampal structure, the dentate gyrus.

**Figure 4:**
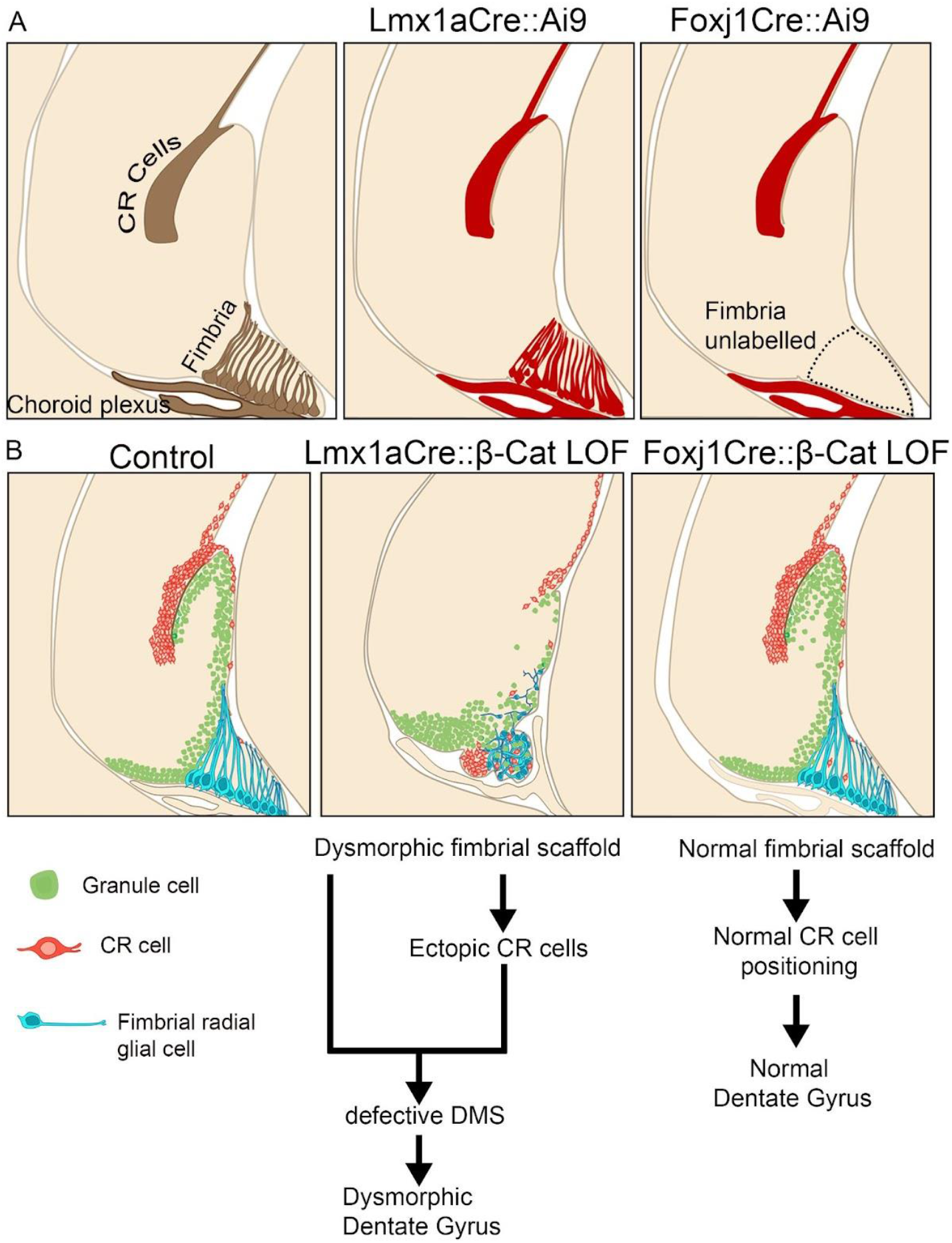
(A) The cartoons illustrate all three hem-derived lineages (brown) i.e., the choroid plexus, CR cells, and the fimbrial scaffold in the E18.5 hippocampus, which also displays the Ai9 reporter (red) when recombined with Lmx1Cre. Foxj1Cre drives recombination in the CR cells and the choroid plexus but not in the fimbrial scaffold. (B) Cartoon of the control hippocampus showing the dentate migratory stream (green), the fimbrial scaffold (blue), and CR cells (orange) in the hippocampal fissure. In the Lmx1aCre::β-Catenin LOF brain fimbrial scaffold is disorganized, resulting in CR cells and dentate migratory cells accumulating in an ectopic protrusion. No such disruption is seen in the Foxj1Cre::β-Catenin LOF hippocampus.

## Materials and methods

### Mice

The Institutional Animal Ethics Committee of the Tata Institute of Fundamental Research Mumbai, India has approved all animal protocols. All mice strains were maintained in an ambient temperature and humidity condition. The food and water were available ad libitum and a 12hr light-dark cycle was strictly followed. The Lmx1aCre mouse line described in this study was a kind gift from Kathy Millen (Center for Integrative Brain Research, Seattle Children’s Research Institute; Lmx1aCre line (Chizhikov & Millen, 2004). Foxj1Cre was a kind gift from Michael J. Holtzman (University of Washington, St. Louis). The β-Catenin exon 2-6 floxed mouse line (Brault et al., 2001) was obtained from Raj Awatramani (Department of Neurology and Center for Genetic Medicine, Northwestern University Feinberg Medical School Chicago, Illinois). The Ai9 reporter mouse line was obtained from JAX labs (Stock No. 007909). Noon of the day when the vaginal plug can be observed was defined as embryonic day 0.5 (E0.5). Only male animals are used for analysis because in the Lmx1aCre animals the transgene is located on the X-Chromosome, and in females, it shows a mosaic expression of the Cre due to random X inactivation. For Foxj1Cre experiments, both male and female embryos were analyzed. All the Controls described in the study were littermates unless and otherwise stated. The following primer sets were used for genotyping: Cre FP: 5’ATTTGCCTGCATTACCGGTC3’, Cre RP: 5’ATCAACGTTTTCTTTTCGG3’, and Cre positive band can be seen at 350bp. For genotyping of the β-Catenin LOF mouse (Exon 2-6 floxed) following combination of primers was used: RM41 5’AAGGTAGAG TGATGAAAGTTGTT3’; RM42 5’CACCATGTCCTCTGTCTATTC3’; RM43 5’TACACTATTGAATCACAGGGACTT3’. This PCR exhibits a wild-type band at 221bp and floxed band at 324bp (Brault et al., 2001).

### Immunohistochemistry

Mouse brain coronal sections were mounted on plus slides (Catalogue number: EMS 71869-11) and dried for 2-3 hours on a slide drier. Slides were then transferred to a slide mailer (Catalogue number: EMS 71549-08) and washed with a solution containing PBS+0.01% TritonX-100 for 10 minutes. Further, the slides were washed with PBS+0.03% TritonX-100 for 5 minutes (2 times). Antigen retrieval was performed in a water bath. In this step, a 10mM sodium citrate buffer having pH=6 was used and the slides were incubated at 900C for 10 minutes inside the water bath. The slide mailer was left for 20 minutes at room temperature for cooling and further washed with PBS+0.01% TritonX-100 for 5 minutes (2 times). A blocking solution containing 5% horse serum in PBS + 0.3% TritonX-100 was added and slides are incubated for 1 hour in a humidified box. This step is followed by overnight primary antibody incubation at 4°C cold room. The next day secondary antibody incubation was done at room temperature for 2 hours in a humidified box covered with aluminum foil. 2 washes with 1x PBS were performed on a rocker followed by incubation with DAPI (Invitrogen catalogue # 62248). Slides were mounted using fluoroshield mounting media (Sigma catalogue # F6182) and imaged in an Olympus FluoView 1200 confocal microscope. The primary antibodies used were: LEF1 (Rabbit, 1:200 CST catalogue # C12A5), β-CATENIN (Mouse 1:200, BDbiosciences catalogue # 610153), β-CATENIN (Rabbit 1:50, CST catalogue # 8814), RFP (Rabbit, 1:200 Abcam catalogue # ab62341), RFP (Mouse, 1:200 Invitrogen catalogue # MA5-15257), TBR2 (Rabbit, 1:200 Abcam catalogue # ab23345), TBR2 (Rat, 1:200 Invitrogen catalogue # 14-4875-82), PAX6 (Rabbit, 1:500 Abcam catalogue # ab195045), PROX1 (Rabbit, 1:500 Millipore catalogue # ab5475), BLBP (Rabbit, 1:200 sigma catalogue # ABN14),TRP73 (Rabbit, 1:200 CST catalogue # 14620S), REELIN (Mouse, 1:200 Millipore catalogue # MAb5364), TTR (Rabbit, 1:75 Dako catalogue # A0002), GFAP (Rabbit, 1:200 Sigma catalogue # G9269), GFAP (Mouse, 1:200 Sigma catalogue # G3893), N-CADHERIN (Mouse, 1:200 BDbiosciences catalogue # 610920), NEUROD1 (Rabbit, 1:1000 Abcam catalogue # AB213725), Phospho HISTONE H3 (Rabbit, 1:200 CST catalogue # H0412), Ki67 (Rabbit, 1:200 Abcam catalogue # ab15580), SOX9 (Rabbit, 1:200 Abcam catalogue # ab185230), SOX2 (Mouse, 1:200 Invitrogen catalogue #MA1-014), NFIA (Rabbit, 1:500 Abcam catalogue # ab228897), NFIB (Rabbit, 1:500 Abcam catalogue # ab186738), ALDH1L1(Rabbit, 1:200 Abcam catalogue #ab87117), AQP4 (Rabbit, 1:200 CST catalogue # 59678). Secondary antibodies used in this study are: Goat Anti Rabbit Alexa fluor 488 (1:200, Invitrogen catalogue # A11034), Goat Anti mouse Alexa fluor 594 (1:200, Invitrogen catalogue # R37121), Goat Anti Rabbit Alexa fluor 568 (1:200, Invitrogen catalogue # A11011), Donkey Anti rabbit Alexa fluor 647 (1:200, Invitrogen catalogue # A31573), Goat Anti Mouse Alexa fluor 647 (1:200, Invitrogen catalogue # A21236), Goat Anti Mouse Alexa fluor 488 (1:200, Invitrogen catalogue # A28175), Goat Anti Rat Alexa fluor 488 (1:200, Invitrogen catalogue # A11006)

### In situ hybridization

Plasmids used to generate probes for in situ hybridization were kind gifts from Elizabeth Grove (*Lmx1a, Wnt3a, Axin2*), and Cliff Ragsdale (*Fzd1*,). The detailed procedure of in situ hybridization was previously described in (Tole & Patterson., 1995, Parichha et al., 2022).

### Image acquisition and analysis

Bright-field images were acquired using Zeiss Axioskop-2 plus microscope combined with a Nikon DS-fi2 camera and NIS Elements V4.0 software. Mouse sections were imaged in Olympus FluoView 1200 confocal microscope with FluoView software. All the image analysis was done on Fiji-ImageJ, and/or Adobe Photoshop CS6. 3D reconstruction (Fig. S4, Supplementary movie 1) was performed in Imaris (V7.2.3). In Fig. 3A and H, image stitching was performed using the “pairwise stitching” plugin in Fiji. For all stitching operations “subpixel accuracy” parameter was selected. Nonlinear operation such as gamma correction was not performed in any of the figures. Brightness and contrast adjustments were performed identically for control and mutant conditions. For Fig. 2E and F, cell counting was performed using the “cell counter” plugin in Fiji. Three sections from different rostro caudal levels were scored for each brain. A (317X137μm) ROI was placed along the hippocampal marginal zone of coronal sections to score the cells and was designated as “MZ” (Fig. 2F). Any cells outside that defined ROI is considered as “outside MZ”. 3 sections spread over different rostro caudal levels per brain were quantified. To quantify the NCAD distribution in Fig. 2K and Fig. S5 D & E, a line (3.5um in length) was placed across the cell boundary when the BLBP channel was selected. 10 such ROIs were placed per brain and the intensity profiles were plotted from the NCAD channel using the “multiplot” option in the ROI manager. Every fourth slice of a Z stack was used for quantification. For Fig. 3D TBR2+ cells in the 2 and 3ry are counted using the “cell counter” plugin. The “C” shaped curve of the section where the ventricular zone ends is considered as the beginning of the 2 and 3ry matrix. The “ectopic site” in the Lmx1aCre::β-Catenin LOF brains refers to the prominent blob protruding in the ventricles. 3 sections spread over different rostro caudal levels per brain were quantified. In Fig. S1 quantification of number of PHH3+SOX2+Ai9+ cells (H) and Ki67+SOX2+Ai9+ (I) cells were done using the “cell counter” plugin in Fiji. All the schematics were prepared using Microsoft PowerPoint 2016 and Adobe Photoshop 2017.

### RNA sequencing and analysis

All dissections were performed in ice-cold PBS. For dissecting hem, the telencephalic hemispheres were isolated using #5 forceps under a stereo zoom microscope (Nikon SMZ445). The Choroid plexus was gently removed from the telencephalic ventricle using blunt forceps. Using fine twizzers the hem region was extracted (Fig. S3A). We examined the brains for Ai9 fluorescence before and after microdissection to ascertain that the Ai9-positive hem is properly extracted (Fig. S3A). RNA extraction was performed using a standard Trizol? reagent. Extracted RNA was analyzed for integrity using the Agilent 2100 bioanalyzer (RIN > 7.5). Hem dissected from 8 embryos was pooled for each of two biological replicates. Library preparation and sequencing were performed on the Illumina platform to achieve 100bp or 150bp reads to generate 30 million paired-end reads. The reads were quantified using Feature counts (Liao et al., 2014). Differential expression analysis was performed using DESeq2 on the R platform (v3.6.3).

### Statistics and reproducibility

Biological replicates (n) denote samples obtained from individual embryos/ pups. Blinded quantification was not possible for Lmx1aCre::β-Catenin LOF brains because the mouse genotypes were easily distinguishable by distinct phenotypic features. For Foxj1Cre::β-Catenin LOF brains blinded analysis was performed. While analyzing the images, to avoid bias stringent measures are taken as described in the “Image analysis” section. All statistical analyses were performed in Graph Pad Prism (V9.3.1), Information about the exact statistical test performed and “P-value” information was provided in corresponding figure legends. The distribution of the data points was analyzed using the Kolmogorov-Smirnov normality test. If the data is distributed normally then a parametric test was chosen otherwise a non-parametric test was performed. For all statistical tests, the chosen confidence interval was always 95% (α=0.05). For all violin plots, Solid thick black line represents the median, solid thin black lines represent quartiles, and dots represent individual data points. For all the tests outlier removal was not performed. For the XY plots in Fig. 2K and Fig. S5 error bars (shades) represent SEM. All raw data points are attached in the source data file.

## Supporting information

Contains raw data points for all graphs

file related to RNA seq data in Fig S3

## Data availability

All the raw data are available on request from the corresponding author. The mouse hem E12.5 RNA-seq data (Fig. S3) generated in this study have been deposited in the GEO database under the Bio project id: PRJNA837732 and will be publically available by the time of publication. The excel file comprising the list of differentially expressed genes is attached with this manuscript as supplementary table 1.

## Author contribution

This work was supervised by S. Tole. AP made the first observations of the β-Catenin LOF phenotypes reported in Figure 2. VS & AP performed the mouse RNAseq analysis. AP and DD performed all the mouse immunohistochemistry. MC performed in situ hybridization experiments. MJH provided the Foxj1Cre mouse line. AP, DD, and S.Tole wrote the manuscript and prepared the figures.

## Acknowledgments

We thank Dr. K. Millen (Seattle Children’s hospital) for the kind gift of the Lmx1aCre line, Raj Awatramani (Department of Neurology and Center for Genetic Medicine, Northwestern University Feinberg Medical School Chicago, Illinois) for the β-Catenin Cko line, and Michael J. Holtzman (University of Washington, St. Louis) for the Foxj1Cre line. We thank Dr. Shital Suryavanshi and the animal house staff of the Tata Institute of Fundamental Research (TIFR) for their excellent support. We acknowledge Ms. Binita Vedak for her help in genotyping. This work was supported by a Wellcome Trust-Department of Biotechnology India Alliance Early Career Fellowship (IA-E-12-1-500765, MC); by the Canada-Israel Health Research Initiative, jointly funded by the Canadian Institutes of Health Research, the Israel Science Foundation, the International Development Research Centre, Canada and the Azrieli Foundation (ST, grant # 108875); intramural funds from TIFR-DAE (12-R&D-TFR-5.10-0100RTI2001), and a grant from Department of Science and Technology (DST), Govt. of India (ST; DST/CSRI/2017/202)

## Competing interests

MJH is the Founder and President of NuPeak Therapeutics and a member of the Data Safety Monitoring Board for AstraZeneca. All other authors declare no competing interests.

## Supplementary figures

**Figure S1:**
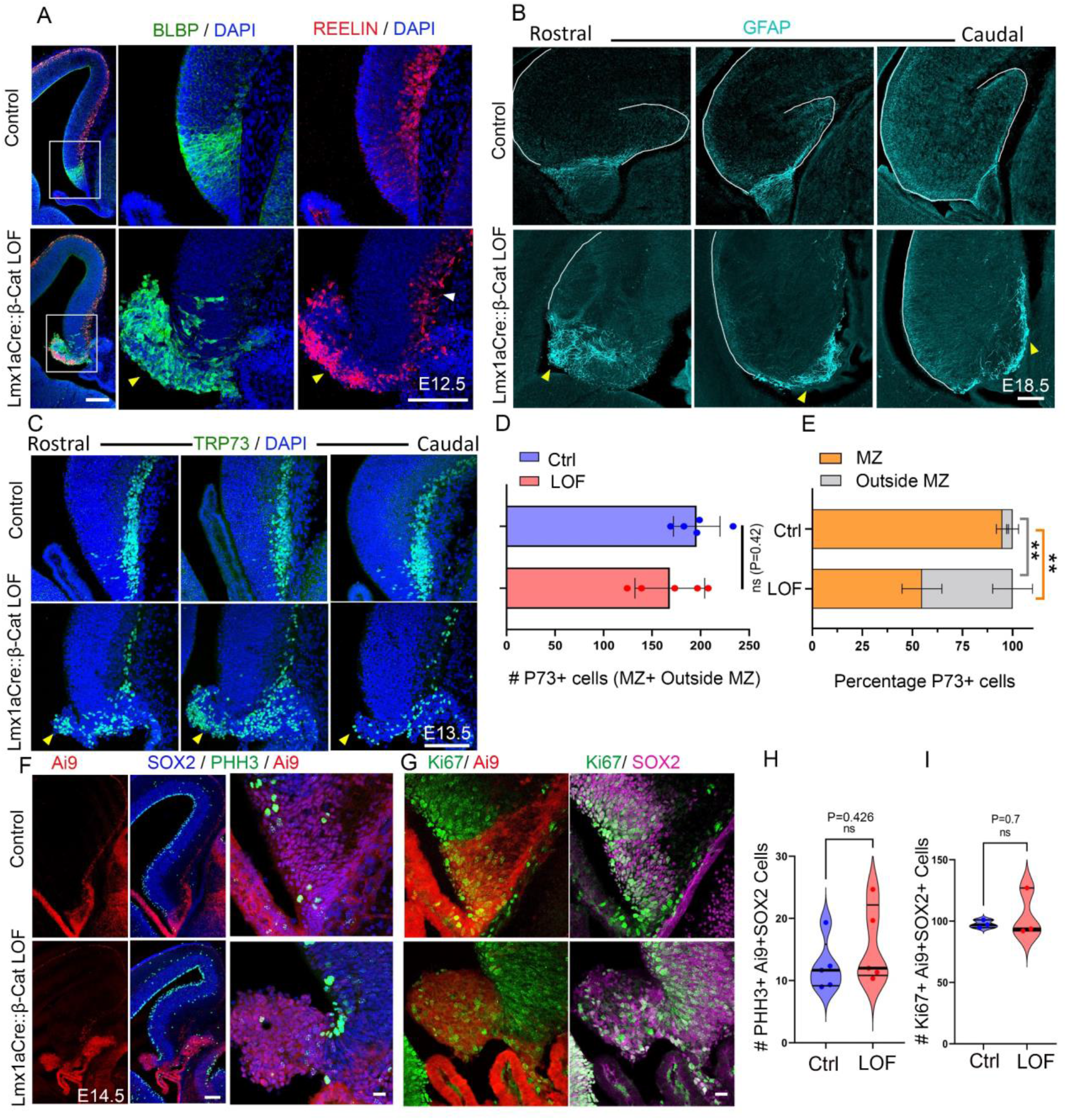
Loss of β-CATENIN in the cortical hem causes dysmorphia of the fimbrial scaffold and mispositioning of the CR cells. (A) Co-immunolabeling for REELIN and BLBP at E12.5 reveals a disorganized fimbrial scaffold and mislocalized CR cells in Lmx1aCre::β-Catenin LOF brains compared with controls. These images show the individual BLBP (green) and REELIN (red) channels of the image in Fig. 2C). (B) GFAP labeling at E18.5 reveals a well-ordered fimbrial scaffold in controls, but a disorganized scaffold in the Lmx1aCre::β-Catenin LOF brains (yellow arrowheads) at multiple rostro caudal levels; N=3 brains (biologically independent replicates) examined over 2 independent experiments. (C) A rostro caudal series reveals that TRP73+ hem-derived CR cells are mislocalized to the ectopically protruding glial mass (yellow arrowheads) throughout the rostro caudal extent of the E13.5 brain. (D) A bar graph representing the total number of TRP73+ cells (Mz+ Outside Mz) shows no significant difference between Lmx1aCre::β-Catenin LOF and control brains. (E) A stacked percentage bar graph displays the altered distribution of TRP73+ cells in Lmx1aCre::β-Catenin LOF compared to controls; N=5 (D and E) brains (biologically independent replicates) examined over 4 independent experiments, bars represent mean±SEM. (F, G) Co-immunostaining for proliferation markers PHH3 and Ki67, and progenitor marker SOX2 (H, I) Violin plots showing quantification of number of PHH3+SOX2+Ai9+ cells (H) and Ki67+SOX2+Ai9+ cells (I) reveal no change in proliferation in Lmx1aCre::β-Catenin LOF brains. Statistical tests (D, E, H and I): Two-tailed unpaired Mann Whitney U test, * p < 0.05, ** p < 0.01, *** p < 0.001, ns if p-value> 0.05; P=0.42 (D), P=0.0079 (E), P=0.426 (H), and P=0.7 (I). All scale bars: 100 μm.

**Figure S2:**
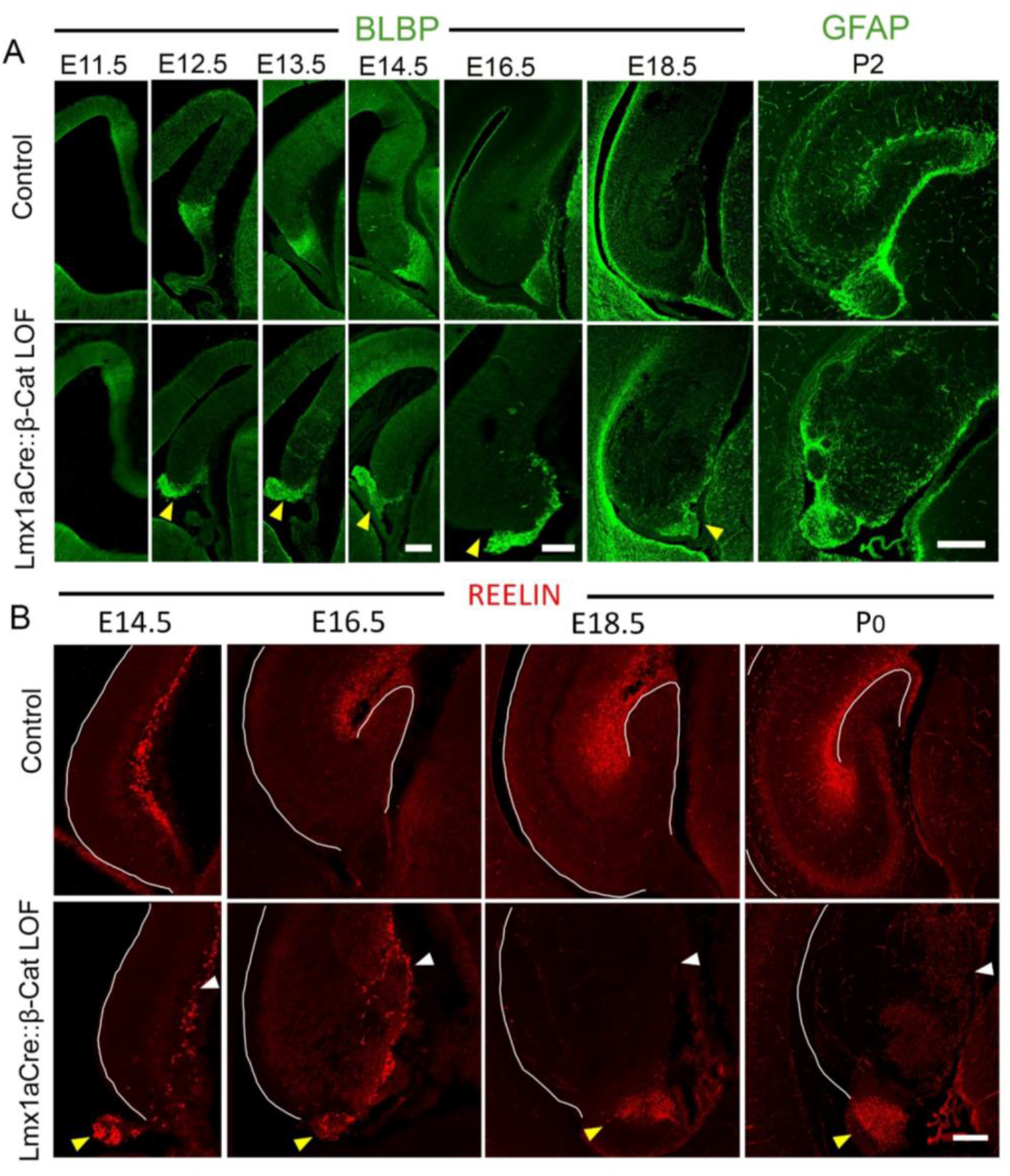
No improvement of the β-Catenin LOF fimbrial scaffold disruption or CR cell mispositioning during late embryonic development. (A) BLBP (E11.5-E18.5) and GFAP (P2) immunolabeling show a well-organized fimbrial scaffold in controls, but a misoriented and disorganized scaffold in the Lmx1aCre::β-Catenin LOF brains; N=3 (E11.5) N=4 (E12.5); N=5 (E13.5 and E14.5), N=4 (E16.5 and E18.5), N=3 (P2) brains (biologically independent replicates) examined over 2 independent experiments. (B) Immunostaining for REELIN at E14.5, E16.5, E18.5 & P0 reveals no improvement in the mislocalized CR cells in the Lmx1aCre::β-Catenin LOF brains during development (Marginal zone: white arrowheads, ectopic localization: yellow arrowheads); N=3 (E14.5); N=5 (E16.5), N=3 (E18.5), N=3 (P0) brains (biologically independent replicates) examined over 3 independent experiments. All scale bars: 100 μm.

**Figure S3:**
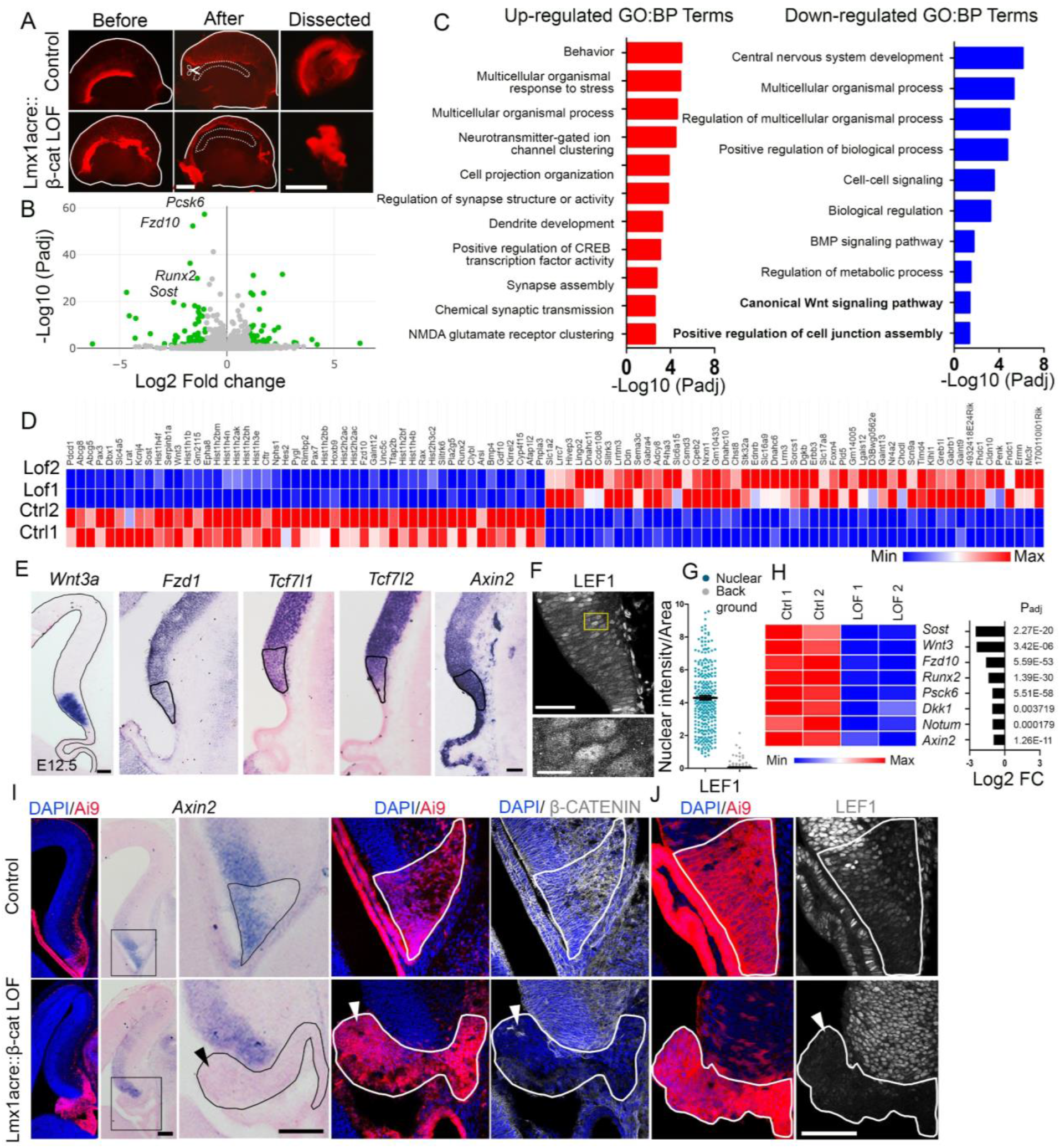
Transcriptional changes associated with loss of β-CATENIN in the cortical hem. (A) Microdissection of E12.5 Ai9-labeled telencephalic hemispheres (before and after dissection) to obtain hem tissue for RNA sequencing. (B) A volcano plot showing differentially regulated genes (green dots represent genes with significantly changed expression). (C) Top 10 upregulated and downregulated GO: BP terms show down-regulation of “Canonical Wnt signaling pathway” and “Positive regulation of cell junction assembly” in Lmx1aCre::β-Catenin LOF compared to controls. (D) Heat map representing Top 50 upregulated (red) and downregulated (blue) genes in control and Lmx1aCre::β-Catenin LOF brains; N=2 biological replicates. (E) The cortical hem, identified by *Wnt3a* expression, expresses several canonical Wnt pathway components such as *Fzd1, Tcf7l1, Tcf7l2*, and *Axin2* at E12.5, N=3 brains (biologically independent replicates) examined over 3 independent experiments. (F, G) immunostaining for Wnt target LEF1 and quantification. (H) Heatmap (displaying normalized reads) and Bar plots (showing log2 fold changes) of candidate Wnt pathway genes downregulated upon loss of β-CATENIN in the hem (complete data in supplementary table 1). (I) *In situ* hybridization for *Axin2* and immunolabeling for β-CATENIN in serial sections shows *Axin2* is undetectable (black arrowhead) at the same region where β-CATENIN staining is lost (Ai9 + area) in Lmx1aCre::β-Catenin LOF brains compared to controls. (J) LEF1 immunolabeling is almost undetectable (white outline) in Lmx1aCre::β-Catenin LOF hem (Ai9+ area). All scale bars: 100 μm.

**Figure S4:**
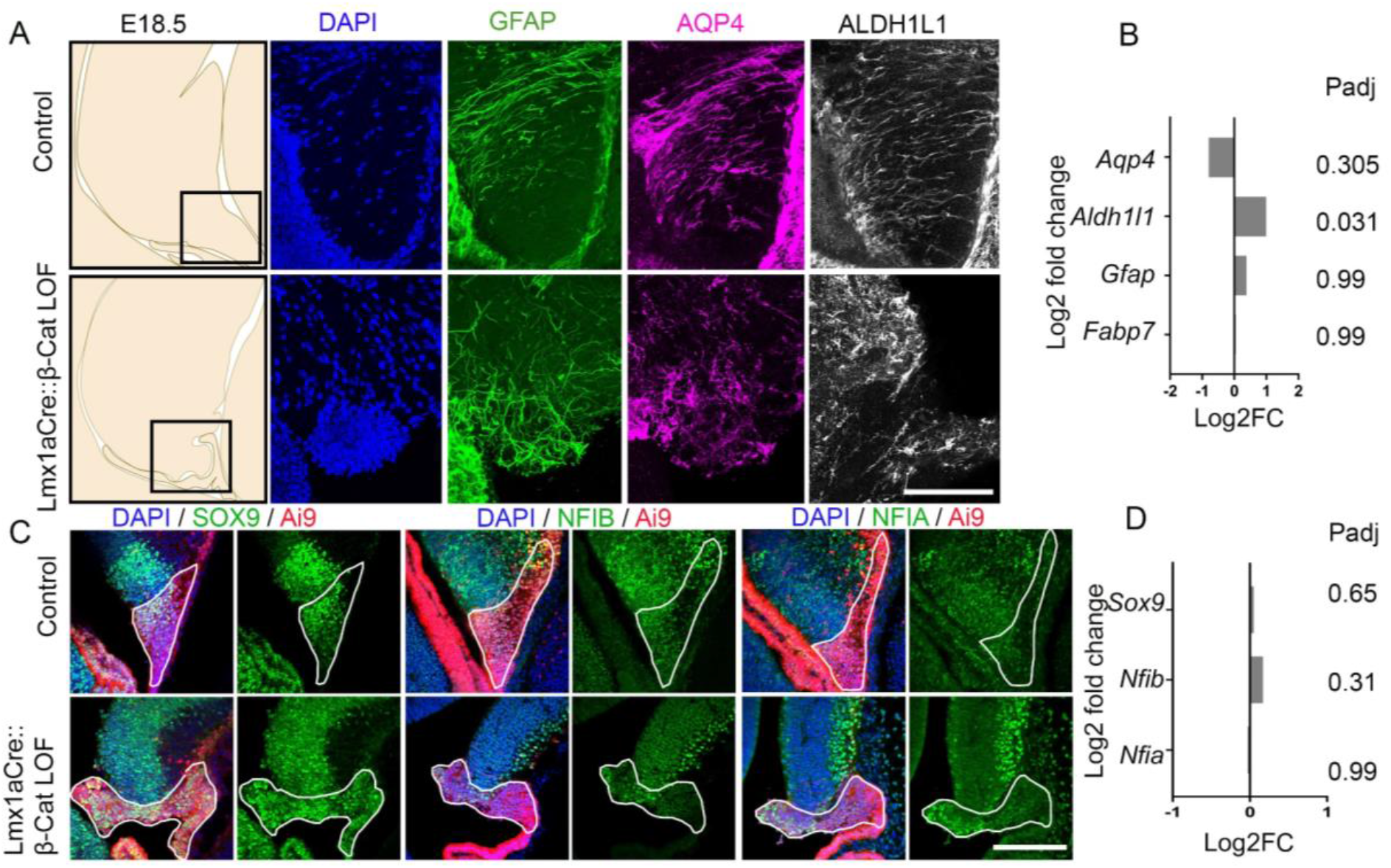
Fimbrial scaffold markers and regulators maintain their expression upon loss of β-CATENIN. (A, B) The β-Catenin LOF fimbrial scaffold continues to display GFAP, AQP4, and ALDHL1 immunoreactivity (A), and expression of *Aqp4, Aldh1l1, Gfap, and Fabp7* is not significantly different (B) from that in control brains. (C) Immunostaining for SOX9, NFIA, and NFIB displays the presence of these regulators of fimbrial scaffold development and (D) Expression of *Sox9, Nfib, and Nfia* is not significantly altered in both control and β-Catenin LOF brains.

**Supplementary Figure S5:**
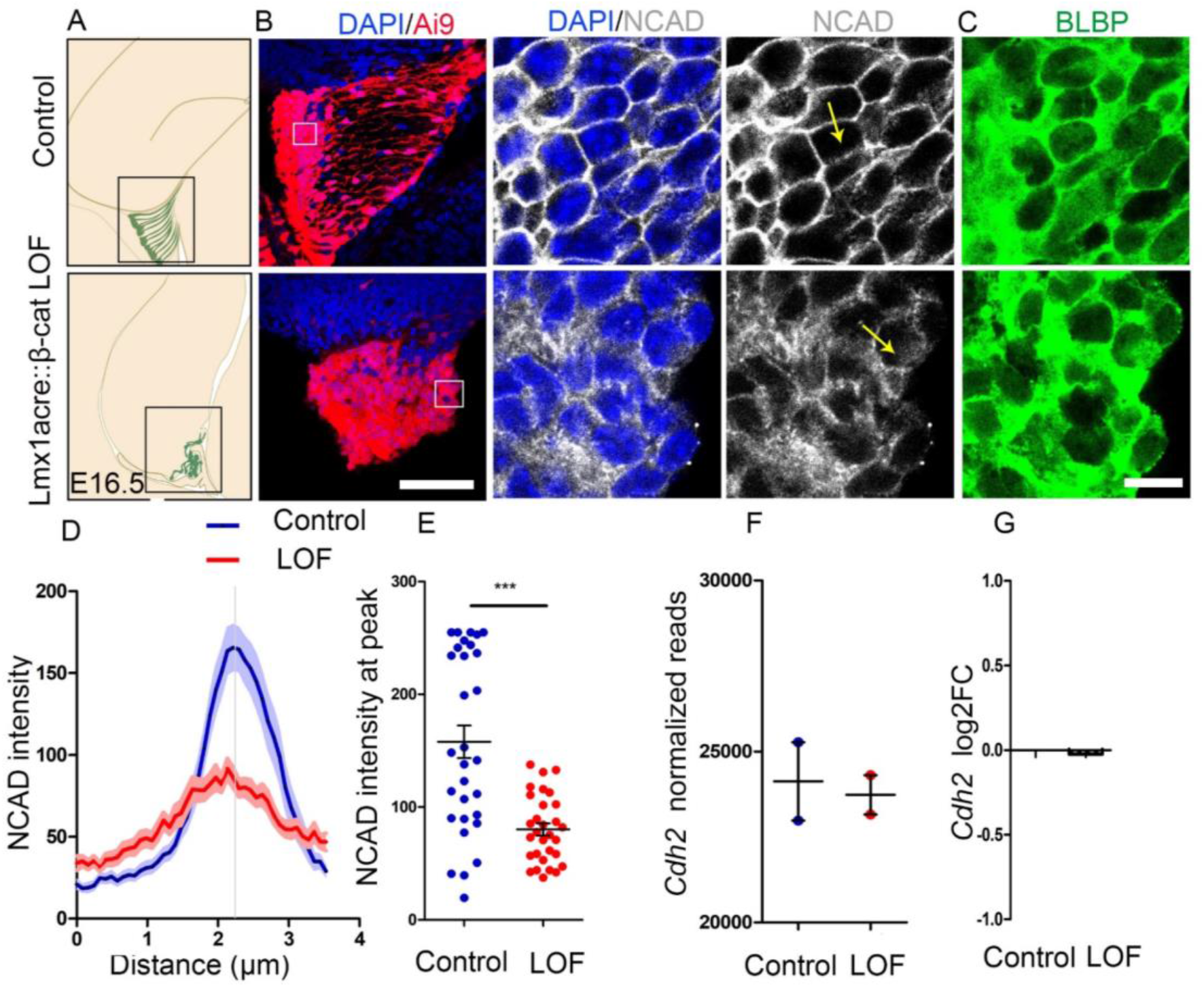
Loss of β-CATENIN in cortical hem leads to altered cell-cell adhesion. (A-C) Distribution of adherens junction marker N-CADHERIN and fimbrial scaffold marker BLBP in Ai9+ hem-derived cells. (B) High magnification images correspond to the boxed regions in the Ai9 image. (D, E) The intensity distribution of N-CADHERIN was quantitated across cell junctions (along the yellow arrow in the NCAD image). N= 3 brains (biologically independent replicates) examined over 3 independent experiments. (G) Normalized read counts (F) and log2 fold change for *Cdh2* gene, which encodes for NCAD, shows no significant change (p=0.99) between Lmx1aCre::β-Catenin LOF brains compared to controls. Statistical test: Two-tailed unpaired t-test with Welch correction (E) and Two-tailed unpaired Mann Whitney U test (D), * p < 0.05, ** p < 0.01, *** p < 0.001, ns if p-value> 0.05; P<0.0001 (E). Scale bars: 10 μm (C); 100 μm (B).

**Supplementary Figure S6:**
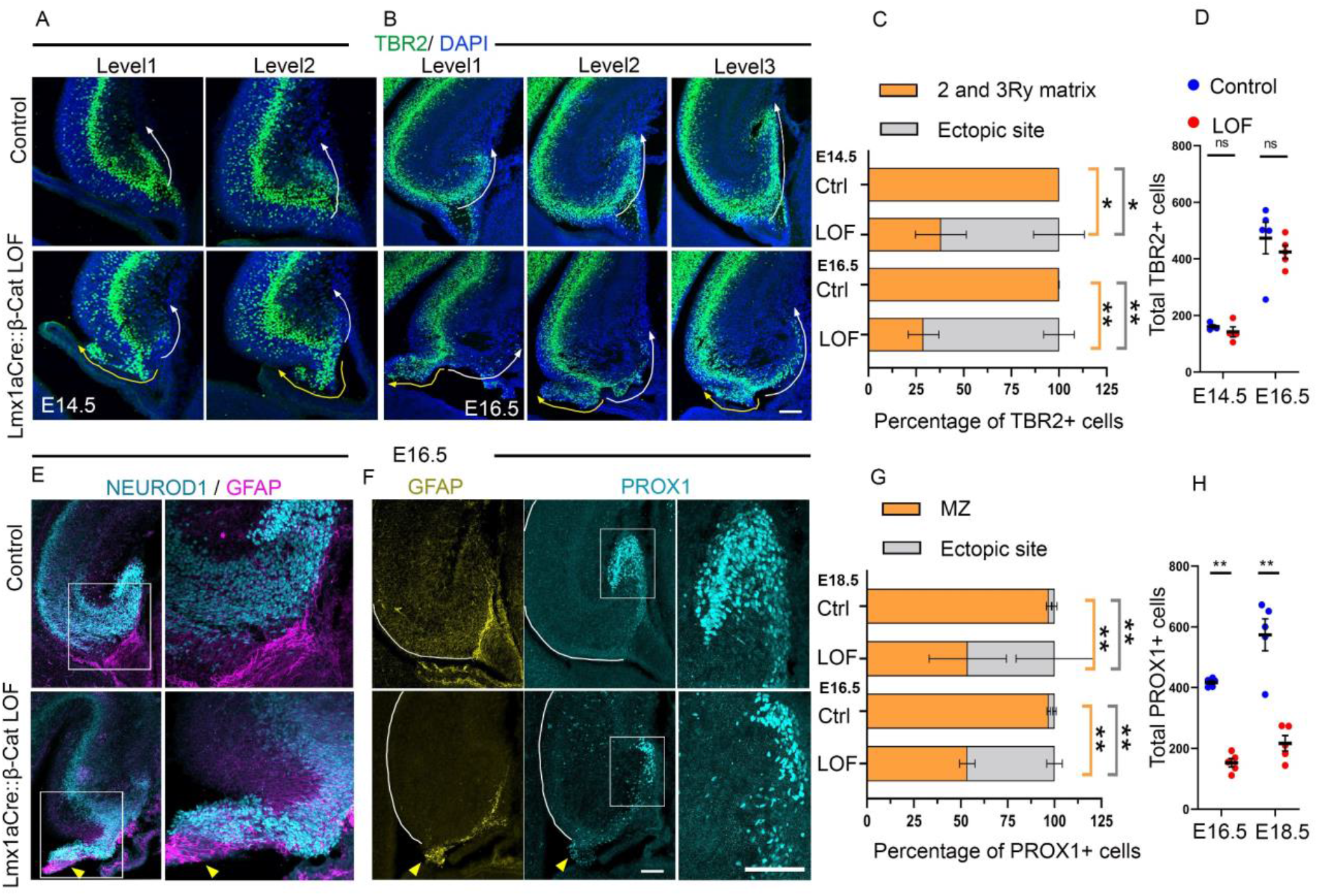
The dentate migratory stream is diverted in the Lmx1aCre::β-Catenin LOF brains. (A, B) Sections at different rostro-caudal levels from E14.5 (A) and E16.5 (B) brains display TBR2+ cells in the 2ry & 3ry matrix of the dentate migratory stream in controls (white arrows). This migration is diverted to the ectopic glial protrusion in Lmx1aCre::β-Catenin LOF brains (yellow arrows). Stacked percentage bar graphs displaying the altered distribution of TBR2+ dentate migratory cells in the 2ry & 3ry matrix vs the ectopic protrusion in Lmx1aCre::β-Catenin LOF brains compared to controls, N=4 (E14.5), N=5 (E16.5) brains (biologically independent replicates) examined over 3 independent experiments. (D) The total number of TBR2+ cells is not significantly different between Lmx1aCre::β-Catenin LOF and control brains. (E, F) At E16.5, Lmx1aCre::β-Catenin LOF brains display a disorganized GFAP+ fimbrial scaffold (yellow arrowheads) and mislocalized NEUROD1+ (E) and PROX1+ (F) dentate migratory cells. The boxed regions in E and F are shown at high magnification in the adjacent images. (G) Stacked percentage bar graphs displaying proportion of PROX1+ granule cells in the MZ (white box in F) vs the ectopic site (outside the white box) in the Lmx1aCre::β-Catenin LOF brains compared to controls, N=5 (E14.5 and E16.5) brains (biologically independent replicates) examined over 3 independent experiments. (H) The total number of PROX1+ granule cells is significantly lower in Lmx1aCre::β-Catenin LOF brains compared to controls. Stacked bar graphs (C and G) use the same data sets shown in the violin plots in Fig 3D and G. Error bars in C, D, G, and H represent SEM. Statistical test: Two-tailed unpaired Mann Whitney U test, * p < 0.05, ** p < 0.01, *** p < 0.001, ns if p-value> 0.05; P=0.028 (C, E14.5), P=0.0079 (C, E16.5), P=(0.3,E14.5; 0.15, E16.5), P= 0.007 (G, E16.5 and e18.5), and P=0.008 (H). All scale bars: 100 μm.

**Supplementary Figure S7:**
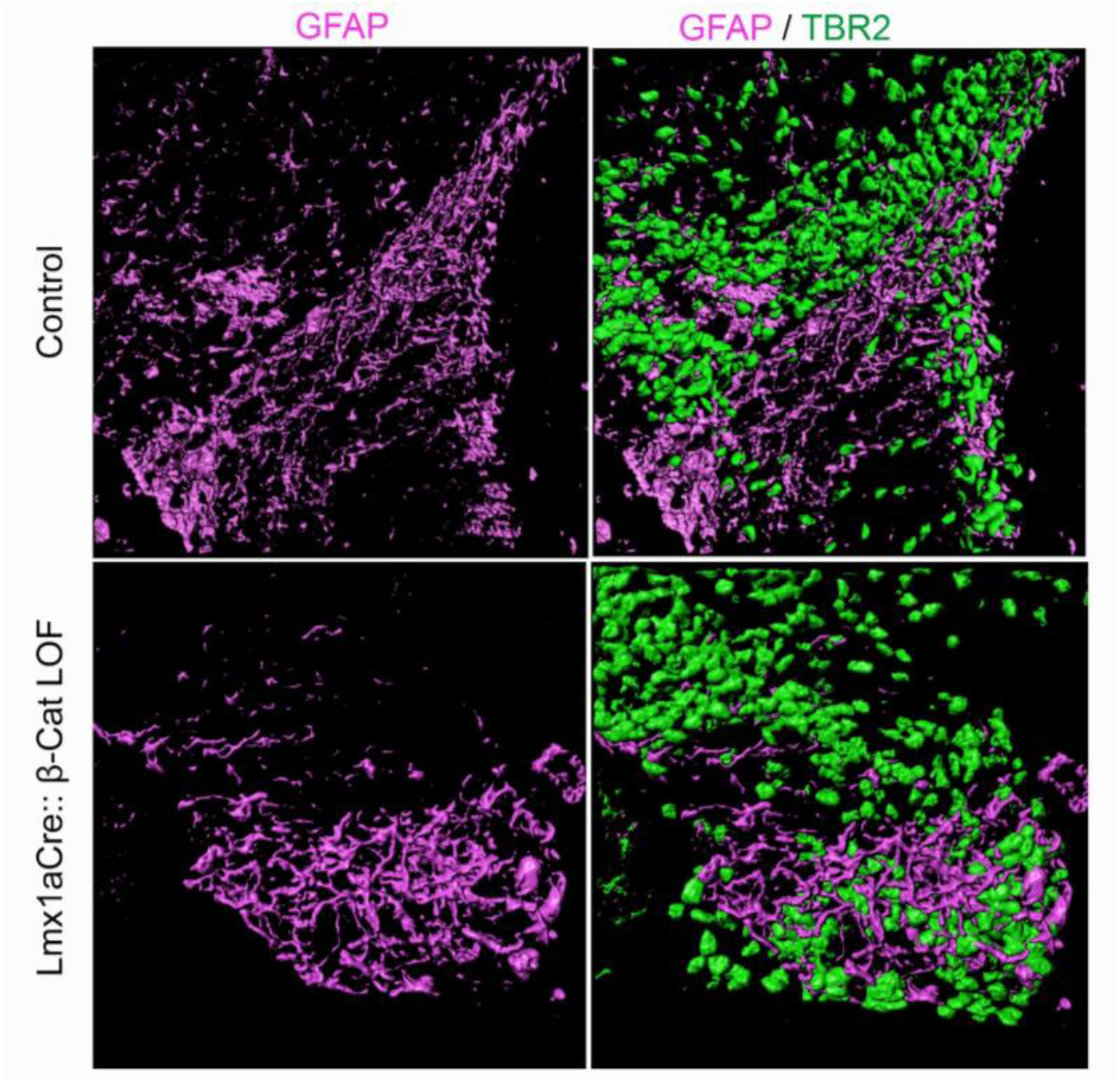
3D reconstruction of the GFAP+ fimbrial scaffold and TBR2+ dentate migratory stream at E16.5. Upon loss of β-CATENIN the fimbrial scaffold is disorganized and the TBR2+ dentate migratory stream is diverted and intermingled with the disoriented fimbrial scaffold (Refer to supplementary movie 1). Scale bar: 30μm.

